# A comparative analysis of the immunotranscriptomic features of DENV-1, -3, and -4 human challenge models

**DOI:** 10.64898/2026.02.17.706422

**Authors:** Céline S. C. Hardy, Lisa A. Ware, Heather Friberg, Joel V. Chua, Kirsten E. Lyke, Stephen J. Thomas, Adam T. Waickman

**Affiliations:** Department of Microbiology and Immunology, State University of New York Upstate Medical University, Syracuse, NY, USA; Global Health Institute, State University of New York Upstate Medical University, Syracuse, NY, USA; Viral Diseases Program, CIDR, Walter Reed Army Institute of Research, Silver Spring, MD, USA; Institute of Human Virology, University of Maryland School of Medicine, Baltimore, MD, USA; Center for Vaccine Development and Global Health, University of Maryland School of Medicine, Baltimore, MD, USA

**Keywords:** Dengue, DENV, DHIM, Immunotranscriptomics

## Abstract

**Background:** Dengue virus (DENV) infections cause a range of clinical symptoms, from a mild febrile illness to severe disease. Higher levels of DENV RNAemia are associated with severe dengue, although this relationship is incompletely understood. Dengue Human Infection Models (DHIMs), in which volunteers are experimentally infected with underattenuated DENV strains, provide an invaluable tool for studying early virologic, transcriptional, and immunologic features of infection. DHIM studies using DENV-1, DENV-3, and DENV-4 have demonstrated qualitatively distinct clinical features, however, the contribution of RNAemia and serotype to divergent transcriptional and clinical profiles in these challenge models remains unclear.

**Methods:** We performed a comparative analysis of DHIM-1, DHIM-3, and DHIM-4 studies to determine shared and unique features of the transcriptional response to infection and their associations with RNAemia and clinical symptoms. We then exposed primary human PBMC *in vitro* to DENV-1 or DENV-3 at varying titers and performed bulk RNA sequencing.

**Findings:** Across DHIMs, we identified a set of conserved, upregulated genes at day of peak RNAemia, representing a core antiviral response independent of serotype. Further, a unique gene signature indicating downregulated cytoplasmic translation emerged in a subset of DHIM-3 participants with elevated RNAemia and symptomatology. *In vitro* PBMC exposure to DENV demonstrated that conserved and unique gene expression signatures varied as a function of viral dose rather than serotype.

**Interpretation:** These data show that viral burden correlates with transcriptional responses and clinical symptomatology following experimental DENV infection, contributing to our understanding of dengue pathogenesis and immunity.

**Research in Context:** *Evidence for this study:* Dengue virus (DENV) infections are known to elicit a range of clinical symptoms. Previous studies have demonstrated the association of higher DENV RNAemia with more pronounced symptoms and elevated risk for severe disease. Understanding the molecular features associated with RNAemia kinetics may provide insight into disease immunologic and clinical pathogenesis. Three live virus human challenge model studies employing underattenuated DENV-1, DENV-3, and DENV-4 have been conducted which demonstrate variance in clinical and virologic features. These experimental DENV infections serve as a valuable model to study early kinetics of infection and immunotranscriptomic features associated with RNAemia irrespective of serotype.

*Added value of this study:* This study is the first to conduct a head-to-head comparison of the transcriptional features of DHIM-1, -3 and -4 models, and to provide an in-depth analysis of these signatures with respect to RNAemia kinetics. These unique datasets provide a rare opportunity to investigate the longitudinal transcriptional signatures associated with peak symptomatology, serotype and RNAemia kinetics.

*Implications of all the available evidence:* These data support the existence of conserved gene expression features of DENV infection, irrespective of serotype and dependent on RNAemia levels. These transcriptional signatures are relevant for our understanding of early events after DENV infection and their relationship to RNAemia as a correlate of disease severity.

## INTRODUCTION

Dengue represents a globally relevant public health threat with nearly 100 million clinically apparent infections annually^1^. Dengue is caused by infection with one of four immunologically distinct dengue viruses (DENV-1 to -4), referred to as serotypes, which frequently co-circulate in endemic areas. Dengue virus infection outcomes range from asymptomatic to life-threatening severe disease, but the factors governing disease outcomes are incompletely understood ^2,3^. Although sequential heterotypic infection and pre-existing immunity have established associations with severe disease^4–6^, substantial heterogeneity in clinical outcomes is observed even among flavivirus naïve individuals, indicating a role for factors other than pre-existing immunity in shaping disease pathogenesis^7^.

Viral burden is known to influence dengue severity^8–14^. Studies have demonstrated that higher DENV RNAemia is associated with increased symptom severity, laboratory abnormalities and risk of progression to severe dengue in natural infection cohorts^12–14^. Although these associations between symptom severity and elevated RNAemia levels have been described, the ability to assess these relationships longitudinally is complicated by limited access to early infection timepoints in the context of natural infection. Furthermore, co-circulation and potential for infection with any of the four DENV serotypes in endemic areas make it difficult to interpret infection history, an important factor known to impact immunologic responses and disease outcomes^15^. Variable exposure history, uncertainty in timing of infection, and limited access to early infection samples are obstacles to completely understanding the kinetics of immunological, virological, and clinical events after a first DENV infection.

Dengue Human Challenge Models (DHIM) address many of these obstacles by exposing flavivirus-naïve adults to underattenuated DENV strains in a controlled fashion with precisely defined and frequent timing of biologic sampling^16^. These models have improved our understanding of early virologic, transcriptomic, and immunologic features of DENV infection ^17–21^. Three separate DHIM studies employing DENV-1, DENV-3, and DENV-4 strains have been conducted by a consortium involving SUNY Upstate Medical University, University of Maryland, and the U.S. Army^18,19,21^.These studies have each demonstrated reproducible RNAemia, mild to moderate dengue-like symptoms, and robust adaptive immune activation, while also demonstrating qualitative differences in virologic and clinical characteristics across models. However, it remains unclear whether differences across models represent serotype specific biology or are driven by other features such as differences in viral burden. Specifically, the extent to which viral replication drives divergence in host transcriptional responses has not been systematically assessed across DHIMs.

Given the association of RNAemia levels and severe disease seen in the natural infection setting, understanding the contribution of serotype and RNAemia levels to transcriptional profiles in the DHIMs may provide valuable insight into mechanisms contributing to variable clinical outcomes. In this work we leverage data from DHIM-1, DHIM-3 and DHIM-4 studies to define shared and unique transcriptomic signatures across models and to determine their relationship to RNAemia. We observed a conserved anti-viral gene expression signature across models at peak RNAemia, in addition to a unique downregulated protein translation signature in DHIM-3 subjects with the highest peak RNAemia titers. The temporality of these signatures followed RNAemia kinetics across DHIMs, which we additionally interrogated by *in vitro* infection of PBMCs at varying viral titers. Consistent with the human challenge model data, transcriptional profiles of PBMC demonstrated that viral ‘dose’, rather than serotype, predominantly shapes transcriptional responses, providing a framework for understanding virologic and immunologic kinetics in these models.

## METHODS

### Study design

Data for this analysis were obtained from published studies, as previously described^18,19,21^. The DHIM and associated analysis was approved by the institutional review board of SUNY Upstate Medical University, the Department of Defense’s Human Research Protection office, University of Maryland Baltimore and the Walter Reed Army Institute of Research (WRAIR). These studies complied with all relevant ethical regulations. In brief, flavivirus-seronegative, healthy volunteers received a single 0.5 mL subcutaneous inoculation containing one of three underattenuated DENV strains: DENV-1 strain 45AZ5 (9 participants, 6.5 × 10^3^ plaque-forming units (PFU)/mL, NCT04298138), DENV-3 strain CH53489 (9 participants, 1.4 × 10³ PFU/mL, NCT05268302), or DENV-4 strain H-241 (4 participants, 1.9 × 10^2^ PFU/mL, NCT05268302). Variable sampling regimens were employed as previously outlined^18,19,21^, where blood draw, saliva collection, clinical evaluation, and self-reported solicited symptoms were collected through days 0-28. Solicited adverse events and clinical laboratory data were collected for all participants. The sampling collection schedule for whole blood biospecimens from which RNA was sequenced is summarized in **Table 1**. Quantitative DENV–specific RT-PCR for DENV-1, DENV-3, or DENV-4 was previously performed using published techniques^18,19,21^.

**Table 1.** Days of RNAseq sample collection and analysis.

| Study | Subject | Days analyzed | Day of peak RNAemia | Peak RNAemia RNAseq sample |
| --- | --- | --- | --- | --- |
| DHIM1 | 201 | 0, 8, 10, 14, 28 | 11 | 10 |
| DHIM1 | 202 | 0, 8, 10, 14, 28 | 14 | 14 |
| *DHIM1* | 203 | 0, 8, 10, 14, 28 | 11 | 10 |
| *DHIM1* | 204 | 0, 8, 10, 14, 28 | 8 | 8 |
| *DHIM1* | 205 | 0, 8, 10, 14, 28 | 9 | 8 |
| *DHIM1* | 206 | 0, 8, 10, 14, 28 | 10 | 10 |
| *DHIM1* | 207 | 0, 8, 10, 14, 28 | 9 | 8 |
| *DHIM1* | 208 | 0, 8, 10, 14, 28 | 11 | 10 |
| *DHIM1* | 209 | 0, 8, 10, 14, 28 | 11 | 10 |
| *DHIM3* | 301 | 0, 6, 8, 10, 14, 28 | 6 | 6 |
| *DHIM3* | 302 | 0, 6, 8, 10, 15, 28 | 8 | 8 |
| *DHIM3* | 303 | 0, 6, 8, 9, 16, 28 | 8 | 8 |
| *DHIM3* | 304 | 0, 6, 8, 10, 15, 28 | 7 | 8 |
| *DHIM3* | 305 | 0, 6, 7, 9, 15, 28 | 6 | 6 |
| *DHIM3* | 306 | 0, 6, 8, 10, 14, 28 | 6 | 6 |
| *DHIM3* | 307 | 0, 6, 8, 10, 14, 28 | 8 | 8 |
| *DHIM3* | 308 | 0, 6, 8, 9, 13, 28 | 6 | 6 |
| *DHIM3* | 309 | 0, 6, 8, 12, 14, 28 | 6 | 6 |
| *DHIM4* | 4021 | 0, 2, 4, 6, 8, 10, 12, 14, 28 | 6 | 6 |
| *DHIM4* | 4022 | 0, 2, 4, 6, 8, 10, 12, 14, 28 | 6 | 6 |
| *DHIM4* | 4028 | 0, 2, 4, 6, 8, 10, 12, 14, 28 | 7 | 6 |
| *DHIM4* | 4029 | 0, 2, 4, 6, 8, 10, 12, 14, 28 | 6 | 6 |

Clinical data were collected as previously described^18,19,21^. Briefly, volunteers were monitored for symptoms in an outpatient setting unless hospitalization criteria were met. Solicited adverse events (clinical or laboratory parameters) included fever (≥38°C), rash, headache, muscle pain, joint pain, fatigue, eye pain, bone pain, abdominal pain, nausea and/or vomiting, elevated liver enzymes (alanine aminotransferase [ALT] and aspartate aminotransferase [AST]), leukopenia, and thrombocytopenia. Symptoms were graded as mild, moderate, or severe according to the FDA toxicity scale.

### DHIM RNAseq library preparation, sequencing and gene expression analysis

RNA sequencing for all DHIM samples was performed as previously described^18,19,21^. In brief, whole blood collection using PAXgene RNA collection tubes (Becton, Dickinson Biosciences; Franklin Lakes, NJ, USA) and RNA recovery using the Qiagen PAXgene Blood RNA isolation kit were performed. Sequencing was performed on a 300 cycle Novaseq 6000 (DHIM-1) or Novaseq X (DHIM-3, DHIM-4) instrument using v1.5S4 reagent set.

Gene expression analysis was performed as previously described^18,19,21^. In summary, raw reads were mapped to the human transcriptome (Ensembl, *Homo Sapiens*, GRCh38, ftp.ensembl.org) using Kallisto version 0.46.2. Transcript level counts and abundance data were imported and summarized in R (Version 4.5.0) using the TxImport package (version 1.36.1) and trimmed mean of M values (TMM) normalized using the EdgeR package (version 4.6.3). Log_2_ transformation, normalization, and filtering of data were performed separately for each DHIM analysis involving only peak RNAemia and baseline samples, and analysis involving samples from all timepoints. In the filtering step, genes with more than one Counts per Million (CPM) in at least three samples were included. Example graphs of log_2_ transformation, normalization, and filtering of data are provided in **Figure S1**. In differential gene expression (DEG) analysis, linear modeling (mean variance) and Bayesian statistics were used in the R package Limma (version 3.64.3). DEGs with a log_2_ fold change >1 and *P*-value <0.01 and were considered statistically significant. Plotting of overlapping gene sets was performed using the R package Eulerr (version 7.0.4). In Gene Ontology (GO) pathway enrichment analysis, pathways with a false discovery rate (FDR) corrected p-value < 0.05 were considered statistically significant.

Transcriptomic scores were calculated by summing the TMM normalized abundance (transcript per million (TPM)) of the genes in the indicated gene set (conserved gene sets, unique up or downregulated gene sets). Temporal modelling of transcriptomic scores was performed using R package ggplot2 package (version 4.0.0) geom_smooth() command to compute and visualize splines for each dataset, with specifications method = "gam" and formula = y ∼ s(x, k = 5,bs = "cs") for DHIM data analysis.

### *In vitro* PBMC culture and infection

Whole blood was collected from a healthy human donor, from which PBMC were freshly isolated and plated at 100,000 PBMC per well. PBMCs were inoculated with DENV-1 45AZ5 or DENV-3 CH53489 at 0 (uninfected control), 1 X 10^2^, 10^3^, or 10^4^ PFU (triplicate for each condition). Cells were incubated at 37°C, 5% CO_2_ for 24 hours, following which cells were harvested, washed, and resuspended in RLT plus buffer with β-mercaptoethanol (BMF) and supernatants were discarded.

### PBMC RNAseq library preparation, sequencing, and gene expression analysis

RNA was extracted using Qiagen RNAeasy prep for RNA sequencing. All samples were sequenced on the Illumina NextSeq 2000 using the P3 Reagent kit (300 cycles). Gene expression analysis was performed as described above. Log_2_ transformation, normalization, and filtering of RNAseq data from *in vitro* PBMC infection samples are provided in **Figure S2**. Temporal modelling of transcriptomic scores for *in vitro* data analysis was performed using R package ggplot2 package (version 4.0.0) geom_smooth() command to compute and visualize splines for each dataset, with specifications method = “loess”.

### Statistical analysis

All gene expression analysis was performed in R version 4.5.0. All other analysis was performed using GraphPad Prism (version 10.3.0, GraphPad Software, La Jolla, CA).

## RESULTS

### Clinical and virologic characteristics of DHIM-1, -3, and -4

We first sought to characterize and compare the baseline clinical and virologic characteristics of each DHIM. Each study was structured to incorporate regular sampling between day 0 (day of inoculation) and day 28, with collection of clinical laboratory values, solicited adverse events, PCR for DENV RNA, and select samples sent for bulk RNA sequencing (**Figure 1A**, **Table 1**). **Figure 1B** shows the kinetics of DENV-1, DENV-3 or DENV-4 RNAemia in plasma across the three DHIM models. Duration of RNAemia, average day of peak RNAemia and mean value of peak RNAemia varied across models (**Table S1**). DHIM-3 and -4 demonstrated an earlier peak (day 7 and day 6, respectively) compared to DHIM-1 (day 10) (**Figure 1B**). Elicited symptomatology varied slightly across models, but timing of peak RNAemia aligned with timing of aggregate peak symptomatology in all three DHIMs (**Figure 1C**). The exception to this observation was timing of peak rash symptomatology, which tended to lag slightly behind timing of peak RNAemia. This finding, however, has been previously elaborated such as in the DHIM-1 study where it was found that rash timing was correlated with NS1 antigenemia^18^. Although 100% of participants in both DHIM-1 and DHIM-4 developed a rash, the grade differed, where DENV-4 induced a whole body (grade 3, >50% body surface area) rash, compared to primarily mild grade 1 or 2 rash elicited by DENV-1. DHIM-3 had an increased number of subjects with leukopenia, neutropenia, and elevated liver enzymes (**Figure 1D, Table S1**), demonstrating increased frequency and severity of elicited symptoms compared to DHIM-1 and DHIM-3 cohorts. Together, these data indicate similar RNAemia kinetics and timing of reported symptoms with RNAemia, although with slight differences in the timing of RNAemia and prevalence and severity of symptoms according to each model.

**Figure 1.**
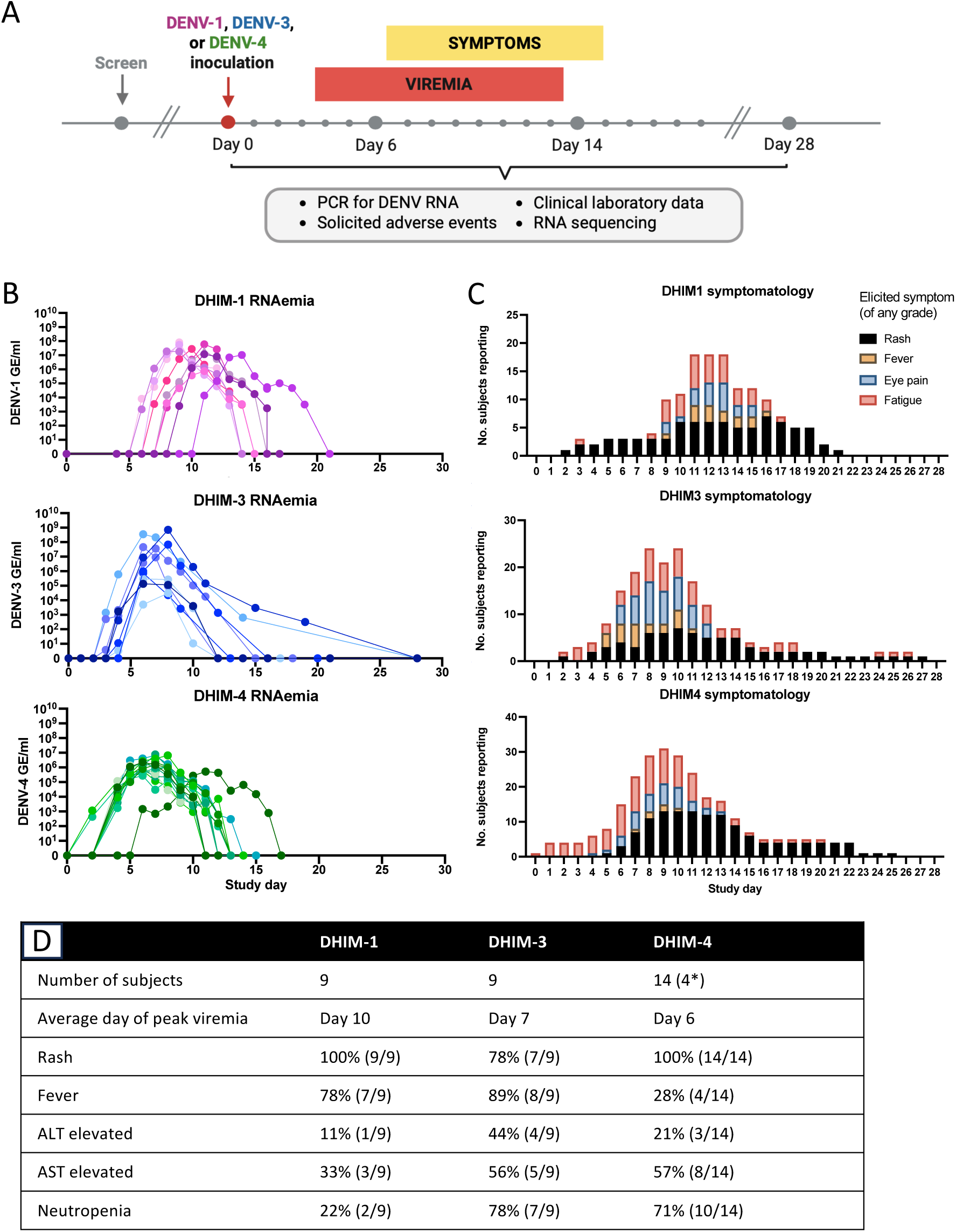
Virologic and clinical features of DENV-1, -3, and -4 human challenge models. **A**) Schematic of study design, timing of sampling and virologic and clinical data collected from DHIM studies. **B**) RNAemia of individual subjects in DHIM-1, -3, and -4 studies in genome equivalents/mL (GE/mL) across study days. Each line is representative of individual study participants. **C)** Distribution of select clinical symptoms (rash, fever, eye pain, and fatigue) according to study day clinical characteristics. **D)** Summary table of key clinical and virologic features of DHIM-1, -3, and -4 studies.

### Conserved and unique gene expression signatures across DHIMs at peak RNAemia

We next aimed to characterize and compare gene expression signatures across models. Given the alignment of reported clinical symptoms with timing of RNAemia, differential gene expression (DEG) analysis was performed at baseline and on the day closest to peak RNAemia for each participant (**Table 1)**. Principal Component Analysis (PCA) comparing day 0 (baseline) to day of peak RNAemia demonstrated global transcriptomic differences of samples (**Figure 2A**). DEG analysis identified many differentially enriched genes from day of peak RNAemia compared to baseline in all models (DHIM-1, n=482; DHIM-3, n=1112, DHIM-4, n=373, **Figure 2B, Tables S2-4**). DHIM-3 had the largest number of DEGs at peak RNAemia compared to baseline and a distinctly large number of significantly downregulated genes (n=277) (**Figure 2B**). Comparison of DEGs enriched at peak RNAemia across DHIM models revealed a common set of 290 upregulated genes (**Table S5**), DHIM-1 and DHIM-4 cohorts had n=46 and n=10 unique DEGs, respectively, and there were n=349 upregulated DEGs unique to DHIM-3 (**Figure 2C, Table S6**). There were no conserved downregulated DEGs across all three models at peak RNAemia. Interestingly, the DHIM-3 cohort demonstrated a unique downregulated gene expression signature (n=266 DEGs) not shared by the other two models (**Figure 2D, Table S6**).

**Figure 2.**
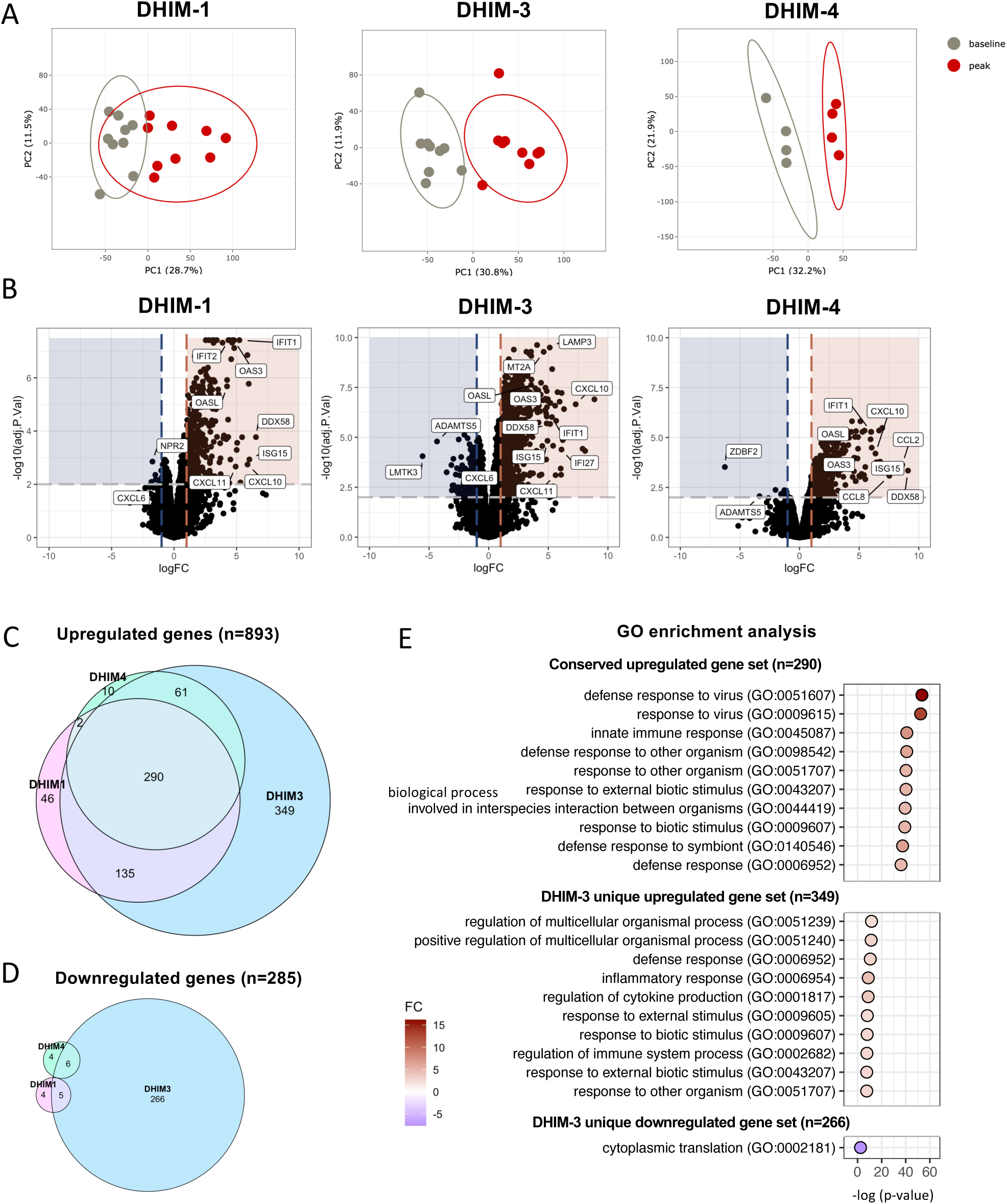
Comparative gene expression analysis at day of peak RNAemia across DHIM-1, - 3, and -4. **A)** PCA analysis across DHIMs comparing transcriptional profiles of individual subjects at day of peak RNAemia (red) compared to baseline (gray). **B)** Volcano plots demonstrating DEGs comparing day of peak RNAemia to baseline. Selected statistically and biologically significant genes are highlighted. Genes with a log_2_ fold change >1 and adjusted p-value < 0.01 were considered significant. **C)** Euler plot demonstrating shared and unique upregulated differentially expressed genes (DEGs) across DHIMs. Pink, DHIM-1; Blue, DHIM-3; Green, DHIM-4 **D)** Euler plot demonstrating shared and unique downregulated differentially expressed genes (DEGs) across DHIMs. Pink, DHIM-1; Blue, DHIM-3; Green, DHIM-4. **E)** Gene Ontology (GO) analysis of DEGs indicating differentially enriched pathways according to conserved upregulated (n=290, top panel), DHIM-3 unique upregulated (n=349, middle panel), or DHIM-3 unique downregulated (n=266, bottom panel) gene sets.

We subsequently performed Gene Ontology (GO) pathway analyses to interrogate the pathways and processes differentially enriched as indicated by these gene sets. GO analysis of the conserved upregulated gene set (n=290) indicated nonspecific activation of various cellular processes in response to virus, indicating a core set of genes with a conserved anti-viral signature across models (**Figure 2E**, top panel). GO analysis of the unique upregulated DHIM-3 gene set indicated similar activation of anti-viral responses, although of lesser magnitude (**Figure 2E**, middle panel). Despite this set of genes emerging as uniquely upregulated in DHIM-3, the similarity of these pathways to those of the conserved gene sets indicates relevant overlap of different genes contributing to the same biologic processes. Interestingly, GO analysis of the DHIM-3 unique downregulated gene set (n=266) indicated a signature of downregulated cytoplasmic translation (GO:002181) (**Figure 2E**, bottom panel). This finding highlights the presence of a unique gene expression profile in DHIM-3 at peak RNAemia. Collectively, these data indicate the existence of both a conserved anti-viral response across DHIMs as well as unique transcriptional features in DHIM-3.

### Divergence of gene expression signatures according to RNAemia level

In the DHIM-3 model, there was a clear divergence in peak RNAemia levels between subjects, with five subjects (subjects 302, 303, 304, 305, 309) experiencing peak RNAemia titers >10^7^ GE/mL and four subjects experiencing peak RNAemia titers <10^7^ GE/mL (**Figure 3A**). No clear divergence in subject RNAemia levels was seen in DHIM-1 or DHIM-4 studies (**Figure 3A**). Higher RNAemia levels in these subjects associated with higher prevalence and severity of symptoms compared to those with lower RNAemia (**Figure 3B**). Although some variability in symptom severity was observed in DHIM-1, the extent of subject-to-subject variation was lower than seen in DHIM-3 and was not accompanied by changes in RNAemia levels^18,19,21^.

**Figure 3.**
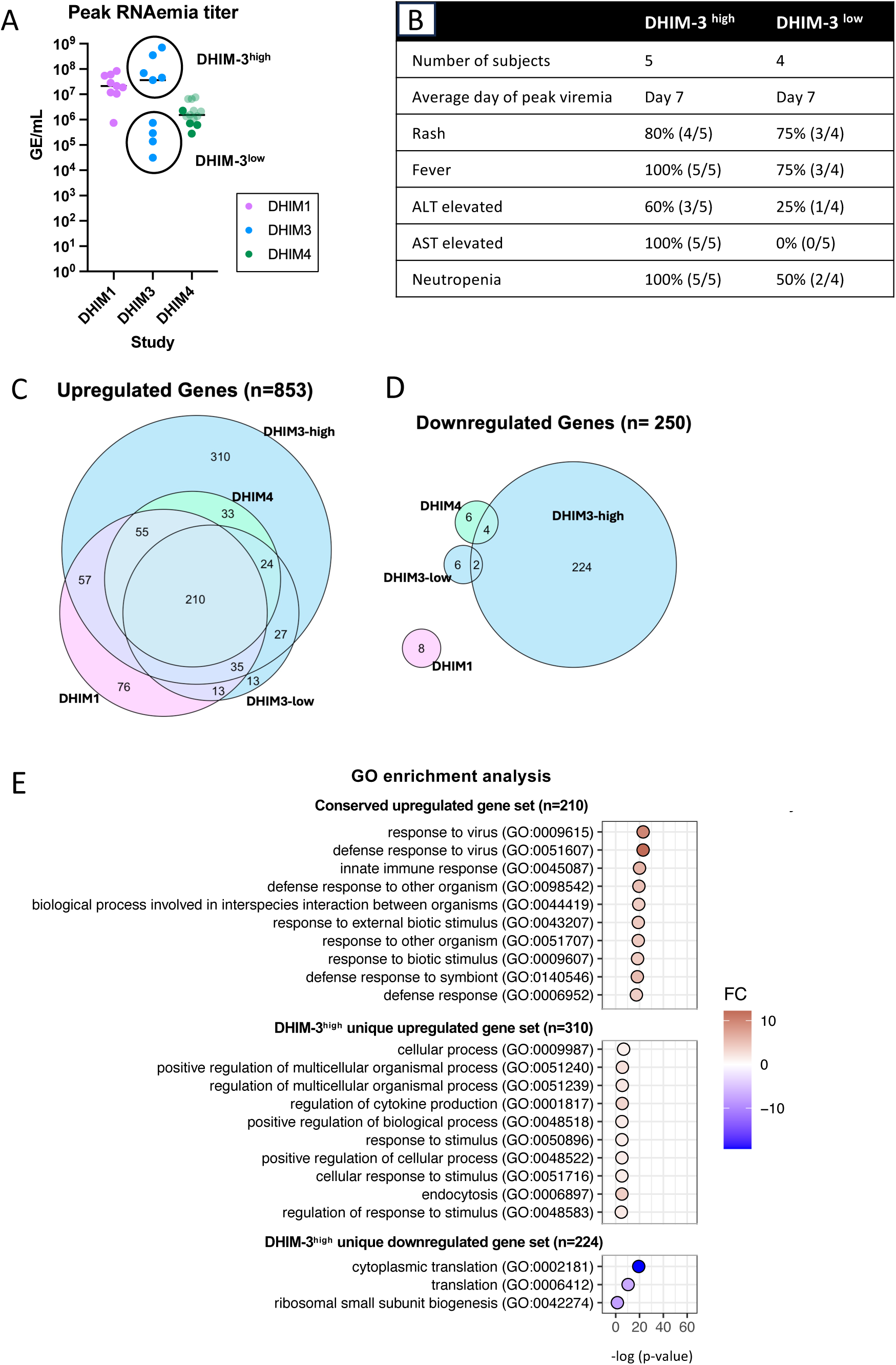
Association of gene expression signatures and RNAemia. **A)** Peak RNAemia of DHIM-1, -3, and -4 subjects, demonstrating bifurcation of high and low RNAemia groups in the DHIM-3 study. **B)** Summary table of key clinical and virologic features comparing DHIM-3^high^ and DHIM-3^low^ RNAemia groups. **C)** Euler plot demonstrating shared and unique upregulated differentially expressed genes (DEGs) across DHIMs. Pink, DHIM-1; Blue, DHIM-3; Green, DHIM-4. **D)** Euler plot demonstrating shared and unique downregulated differentially expressed genes (DEGs) across DHIMs. Pink, DHIM-1; Blue, DHIM-3; Green, DHIM-4. **E)** Gene Ontology (GO) analysis of DEGs indicating differentially enriched pathways according to conserved upregulated (n=210, top panel), DHIM-3^high^ unique upregulated (n=310, middle panel), or DHIM-3^high^ unique downregulated (n=224, bottom panel) gene sets.

Given the previously described association of RNAemia with severe symptomatology in natural infection, we aimed to disentangle the potential contribution of varying peak RNAemia titers to transcriptional signatures that likely drive clinical outcomes. Therefore, we performed a second DEG analysis comparing DHIM-3 participants with high (DHIM-3^high^) versus low (DHIM-3^low^) RNAemia levels to determine whether RNAemia levels drive transcriptional phenotype. DEG analysis comparing signatures at peak RNAemia to baseline for each group (DHIM-1, DHIM-4, DHIM-3^low^ and DHIM-3^high^) was performed (**Table S7-8**) and overlapping DEGs for each group were visualized using Euler plots. When the DHIM-3 cohort was divided into high and low groups, a conserved upregulated gene set still emerged (n=210) (**Figure 3C, Table S9**). However, the unique gene sets from the DHIM-3 study identified in the original DEG analysis emerged as being driven by the DHIM-3^high^ individuals (**Figure 3C-D, Table S10**). These results were consistent at the pathway level, where all key gene sets (conserved upregulated (n=210), unique upregulated DHIM-3^high^ (n=310), and unique downregulated DHIM-3^high^ (n=224)) identified in the four-group analysis indicated similar pathway enrichment to the three-group analysis. Particularly, the downregulated signature in the DHIM-3^high^ group was even more pronounced (**Figure 3E**, bottom panel), with the GO terms “translation (GO:0006412)” and “ribosomal small unit biogenesis GO:0042274)” emerging as additional processes downregulated in the DHIM-3^high^ subjects. These data indicate the existence of a core set of genes upregulated across models, irrespective of peak RNAemia, and a unique signature in DHIM-3 participants with elevated RNAemia and symptomatology.

### Temporality and correlation of RNAemia with gene expression signatures

We next sought to evaluate the expression of these signatures over time to clarify their association with timing of infection and RNAemia. In order to visualize trends in gene set expression across participants over time, transcriptomic scores (TS) were calculated by summing the TMM normalized abundance (transcript per million (TPM)) of the genes in the indicated gene set (conserved gene sets, unique up or downregulated gene sets). Trends of conserved gene set TS expression mimicked patterns of RNAemia across time, where conserved gene set expression increased at time of peak RNAemia but remained low at baseline (day 0) and late time points (day 28) indicating temporal relationships (**Figure 4A, Figure S3**). Additionally, mean peak conserved TS values aligned with mean peak day of RNAemia across studies. The absolute values of these scores differed by study, with DHIM-4 demonstrating lower mean TS than DHIM-1 or DHIM-3 cohorts consistent with the clinical presentation of the models. To further define the relationship of these scores to RNAemia, we performed Spearman correlation of conserved gene set TS with RNAemia (GE/mL) at multiple time points during acute infection. Conserved gene set TS were positively correlated with RNAemia across all models (DHIM-1, R = 0.825, *p* = 6.151e-10; DHIM-3, R = 0.850, *p* = 1.496e-13; DHIM-4, R = 0.826, *p* = 1.844e-08) (**Figure 4B**). When plotted according to the four-group subset (dividing DHIM-3 into high and low RNAemia groups) no differences in trends of the association of RNAemia with TS were notable (**Figure S5**). These associations across DHIM of conserved TS expression indicate that this shared gene expression signature is stable over time, associated with peak RNAemia, and are correlated with RNAemia levels across models.

**Figure 4.**
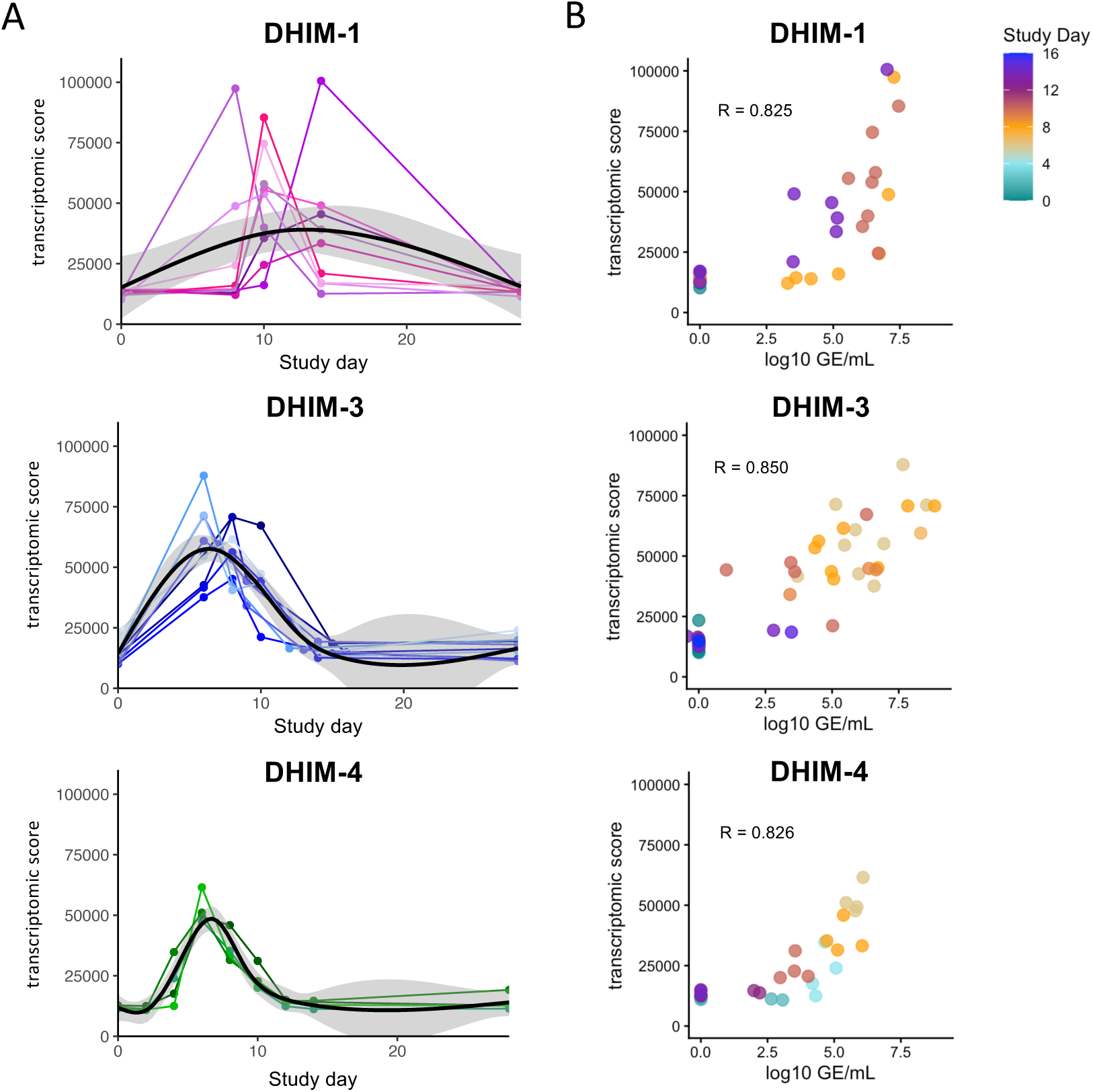
Temporality and correlation of RNAemia and gene expression signatures. **A)** Transcriptomic score (TS) for the conserved upregulated gene set (n=210) across time (study day 0-28) in DHIM-1, DHIM-3 and DHIM-4 subjects. Colored lines and dots indicate individual subjects. A spline-based smoothing curve (black line) illustrates population-level trend between X and Y, with the grey region indicating the 95% confidence interval. **B)** Spearman correlation of RNAemia (log10 GE/mL) and conserved gene set (n=210) TS. Dot color indicates study day according to provided scale.

### Viral titer dictates transcriptional signatures of in vitro PBMC exposure to DENV

Given the observed association of gene expression signatures and RNAemia in the DHIM-3 participants, we then assessed, *in vitro,* whether these signatures were unique to the DENV-3 CH53489 strain. Freshly isolated PBMCs were exposed to DENV at three titers, the highest of which approximated the titer corresponding to the average PFU/mL at peak RNAemia in DHIM participants. PBMC were exposed to no virus (uninfected/mock control) or 1 X 10^2^, 10^3^, or 10^4^ PFU DENV-1 (strain 45AZ5) or DENV-3 (strain CH53489). After 24 hours, cells were harvested and bulk RNA sequencing was performed (**Figure 5A**). PCA analysis showed samples separated primarily along PC2, distinguishing DENV-1 from DENV-3 transcriptional profiles (**Figure 5B**). Further analysis showed the clustering of samples appeared to be primarily driven by the titer of virus exposure and expression of genes in PC1 (**Figure 5C**). DEG analysis comparing each infection condition to mock control also demonstrated increasing numbers of DEGs were associated with increasing titers of virus exposure (**Figure S6**).

**Figure 5.**
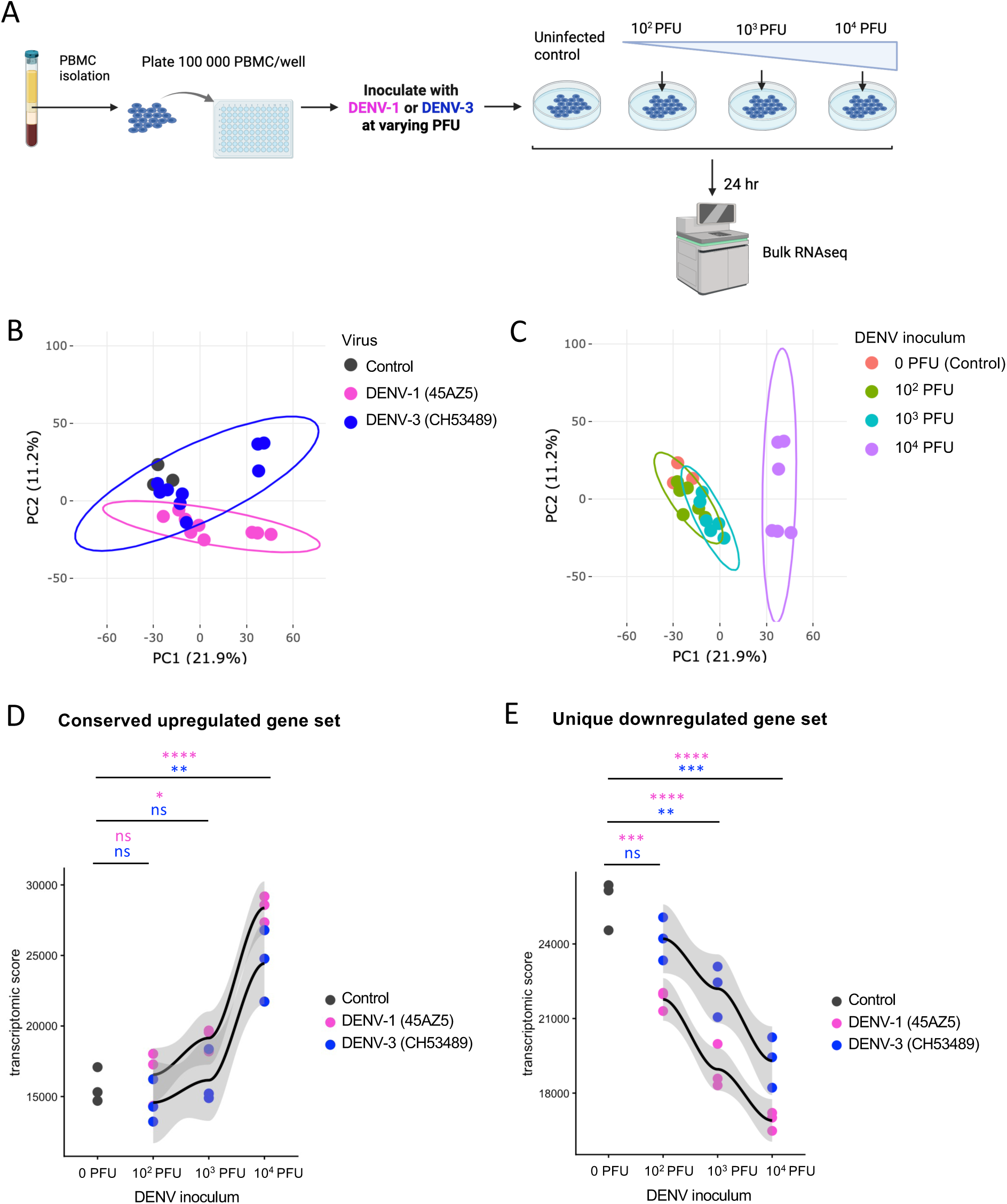
In Vitro PBMC exposure stratifies gene expression according to inoculum dosage. **A)** Schematic representation of experiment design. **B)** PCA plot of gene expression data from *in vitro* exposure of PBMC to DENV-1 or DENV-3. Labelled according to virus used. **C)** PCA plot of gene expression data from *in vitro* exposure of PBMC to DENV-1 or DENV-3. Labelled according to titer of virus used. **D)** Transcriptomic score (TS) of the conserved upregulated gene set (n=210) derived from DHIM analysis. **E)** Transcriptomic score (TS) of the unique DHIM-3^high^ downregulated gene set (n=224) derived from DHIM analysis. Dots represent individual samples. A spline-based smoothing curve (black line) illustrates population-level trend between X and Y, with the grey region indicating the 95% confidence interval.

To assess patterns in expression of the previously defined conserved upregulated (n=210) and unique DHIM-3^high^ downregulated (n=224) gene sets, we calculated corresponding TS values for the *in vitro* samples. Plotted according to the titer of viral inoculum, we observed a trend toward increasing expression of the conserved upregulated gene set (**Figure 5D**). One-way ANOVA with Bartlett’s test (corrected) with Dunnett’s multiple comparisons test indicated a significant, stepwise increase of the conserved upregulated gene set in both DENV-1 (*p* < 0.0001) and DENV-3 (*p* = 0.0007) exposed samples (**Figure 5D**). Likewise, unique downregulated gene expression TS values also indicated a stepwise decrease in expression with increasing virus titer (DENV-1, *p* < 0.0001; DENV-3, *p* < 0.0002) (**Figure 5E**). Together, these *in vitro* data suggest that the transcriptional response of PBMC exposed to DENV-1 or DENV-3 challenge strains is primarily driven by the titer of virus inoculum.

## DISCUSSION

In this study, we compared the virologic, clinical, and immuno-transcriptomic features of the DHIM-1, -3, and -4 experimental human infection cohorts. We observed a conserved set of genes expressed across all three DHIM cohorts corresponding to pathways indicative of immune activation. Our data demonstrated a unique gene expression signature in participants with elevated RNAemia in the DHIM-3 study, indicating downregulated cytoplasmic translation and ribosomal biogenesis. *In vitro* infection of PBMC with the same DENV challenge strains revealed that these signatures increased or decreased accordingly with the titrated level of virus exposure. Together, these data indicate that in these models, the titer of infectious virus exposure drives cellular transcriptomic responses, irrespective of DENV serotype. Our findings provide a framework for understanding the drivers of transcriptional response to DENV infection.

The association between elevated viral burden and altered host transcriptional responses is consistent with prior observations from natural infection^12^. Additionally, multiple studies have demonstrated the association of high RNAemia levels with more severe disease^8–14^. For example, Vuong et al. observed that high RNAemia levels were associated with decreased platelet counts, and that elevated RNAemia levels on each illness day increased the risk of developing severe dengue^14^. However, an important consideration for interpretation of data from many of the studies is that they rely on limited sampling or a single time point, making it difficult to disentangle the temporal relationships of these associations. In this study, the controlled nature of DHIMs allowed the precise alignment of transcriptional changes with virologic kinetics, demonstrating correlation with RNAemia amplitude and highlighting the role of RNAemia as a driving factor of pathogenesis across viral strains.

In this study, we observed a unique downregulated signature of protein translation specific to DHIM-3 participants with elevated RNAemia and symptomatology. Interestingly, a transcriptional signature consistent with downregulated protein translation has been previously described when comparing transcriptional features of severe, natural DENV-1 infection to experimental DENV infection^22^. Although the current study was not powered to formally associate transcriptional signatures with clinical severity, it is notable that these data are consistent with the convergence of elevated RNAemia, increased symptom burden, and translational suppression in a subset of DHIM-3 volunteers. The precise mechanisms by which RNAemia levels are associated with downregulated protein translation and how this mechanism contributes to severe disease are unclear and warrant further investigation.

*In vitro* infection of PBMC in this study further supports a viral-dose-dependent relationship, where exposure of primary PBMC to increasing DENV-1 or DENV-3 titers recapitulated the induction of conserved, upregulated antiviral and downregulated genes. These data suggest the magnitude of viral titer is sufficient to drive these transcriptional programs, and that the DHIM-3 associated signatures we observed are unlikely to be serotype or strain-specific effects.

There are several limitations to this study. All challenge participants were flavivirus naïve at baseline, allowing this study to characterize responses after primary DENV exposure. However, the extension of these findings to the setting of secondary infection where pre-existing DENV or non-DENV flavivirus immunity exists is uncertain. Sample sizes for each DHIM were modest, which may have limited our ability to detect more subtle transcriptional differences, particularly downregulated gene sets in DHIM-1 and DHIM-4 studies. The aggregation of transcriptional analyses across future studies will address this limitation. It is also important to consider that in both the human challenge and *in vitro* experiments, differences in genome-to-PFU ratios of the challenge strain stocks employed complicate direct comparison of infectious dose and assessment of the contribution of noninfectious viral particles. Additional work characterizing the relationship of RNAemia to transcriptional and clinical signatures in response to additional DENV serotypes and strains is of importance. Finally, although DHIM models appear to closely mimic natural DENV infections, it is important to acknowledge there may be immunologic and virologic differences between the two because; 1) challenge viruses are delivered by needle where as wild type DENVs are delivered by mosquito along with potentially immunogenic mosquito salivary proteins, 2) it is unclear what dose of DENV is delivered by a feeding mosquito, 3) challenge DENVs are attenuated by cell passage and chemical mutagenesis, and 4) challenge viruses are delivered to the subcutaneous space while a mosquito delivered virus is mostly likely delivered to a different anatomic space with potential antigen trafficking consequences.

In summary, our study highlights the existence of a conserved gene expression signature across DENV-1, DENV-3, and DENV-4 human infection models. These signatures were enriched at time of peak RNAemia across individuals but additionally correlated with RNAemia kinetics over time. *In vitro*, transcriptional programming also appeared to be primarily driven by viral dose to which human PBMC were exposed. This work contributes to our understanding of the association of transcriptional signatures of inflammation with viremia kinetics. This provides granular and valuable insight into signatures of immunopathogenesis in primary DENV infection. Moreover, these findings are in alignment with previously described associations of RNAemia with severe symptomatology and provide insight into how these kinetics differ across viral serotypes/strains. More broadly, this work informs the interpretation of transcriptional and virologic data for future DHIM studies and natural infection data. These findings support the relevance of viral burden as a unifying determinant of elicited pathogenesis, which may be relevant for advancing our understanding of biomarkers of dengue severity.

## Contributors

**Céline S. C. Hardy**: Conceptualization, Data curation, Visualization, Formal Analysis, Writing – original draft, Writing – review and editing. **Lisa A. Ware**: Methodology, Writing – review and editing. **Heather Friberg**: Writing – review and editing. **Joel V. Chua:** Methodology, Data curation, Writing – review and editing ; **Kirsten E. Lyke**: Methodology, Data curation, Writing – review and editing. **Stephen J. Thomas**: Methodology, Writing – review and editing. **Adam T. Waickman**: Conceptualization, Data curation, Formal analysis, Investigation, Methodology, Funding acquisition, Supervision, Writing – review and editing.

## Acknowledgements

We gratefully acknowledge the technical assistance of Karen Gentile of the Upstate Medical University Molecular Analysis Core (MAC) and the members of the Upstate Global Health Institute (GHI) of SUNY Upstate Medical University. We also wish to thank all the DHIM study participants for making this work possible. Material has been reviewed by the Walter Reed Army Institute. The opinions or assertions contained herein are the private views of the authors and are not to be construed as reflecting the official views of the US Army or the US Department of Defense. Material has been reviewed by the Walter Reed Army Institute of Research. There is no objection to its presentation and/or publication. The investigators have adhered to the policies for protection of human participants as prescribed in AR 70-25.

## Data availability

The authors declare that all data are available within the article, its supporting materials, published data, or public databases. RNA sequencing data for *in vitro* PBMC infection are publicly available through the Gene Expression Omnibus (GEO).

## SUPPLEMENTAL MATERIAL

### Supplemental figures

**Figure S1.** Example log_2_ transformation, normalization, and filtering of bulk RNA sequencing data for all samples from DHIM-1, -3, and -4 studies.

**Figure S2.** Log_2_ transformation, normalization, and filtering of bulk RNA sequencing data for all samples from *in vitro* PBMC exposure to DENV-1 (45AZ5) and DENV-3 (CH53489).

**Figure S3.** Heatmap of conserved upregulated gene set (n=210) expression across study days and cohorts.

**Figure S4.** Heatmap of unique DHIM-3^high^ downregulated gene set (n=224) expression across study days and cohorts.

**Figure S5.** Figure S5. Transcriptomic score expression in DHIM-3^high^ and DHIM-3^low^ groups across study days.

**Figure S6.** PCA analysis and volcano plots of DEG analysis for *in vitro* PBMC DENV-1 (45AZ5) and DENV-3 (CH53489) challenge strain titration.

### Supplemental Tables

**Table S1.** Comparison of virologic and clinical features of DHIM-1, -3 and -4.

**Table S2.** DEGs at day of peak RNAemia compared to baseline in DHIM-1 volunteers.

**Table S3.** DEGs at day of peak RNAemia compared to baseline in DHIM-3 volunteers.

**Table S4.** DEGs at day of peak RNAemia compared to baseline in DHIM-4 volunteers.

**Table S5.** Shared upregulated DEGs at day of peak RNAemia compared to baseline in DHIM-1, DHIM-3, and DHIM-4 cohorts (n=290).

**Table S6.** Unique upregulated (n=349) and downregulated (n=266) DEGs in DHIM-3 volunteers at day of peak RNAemia compared to baseline.

**Table S7.** DEGs at day of peak RNAemia compared to baseline in DHIM-3high volunteers.

**Table S8.** DEGs at day of peak RNAemia compared to baseline in DHIM-3low volunteers.

**Table S9.** Shared upregulated DEGs between DHIM-1, DHIM-3high, DHIM-3low, and DHIM-4 cohorts (n=210).

**Table S10.** Unique upregulated (n=310) and downregulated (n=224) DEGs in DHIM-3high volunteers at day of peak RNAemia compared to baseline.

