## Supplemental figures for "A comparative analysis of the immunotranscriptomic features of DENV-1, -3, and -4 human challenge models"

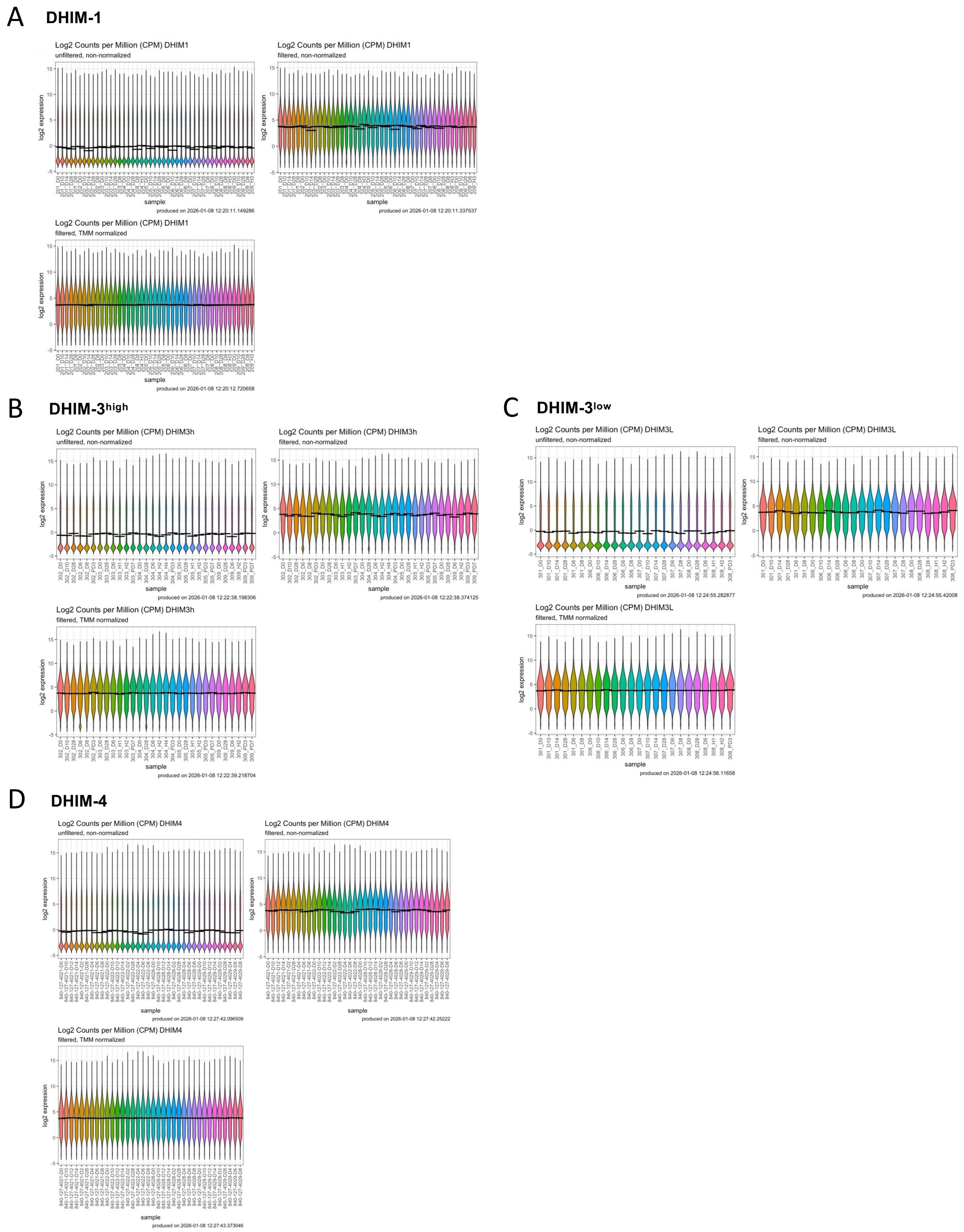


**Figure S1.** Example $\log_2$ transformation, normalization, and filtering of bulk RNA sequencing data for all samples from DHIM-1, -3, and -4 studies. **A)** DHIM-1 samples. **B)** DHIM-$3^{high}$ samples. **C)** DHIM-$3^{low}$ samples. **D)** DHIM-4 samples.

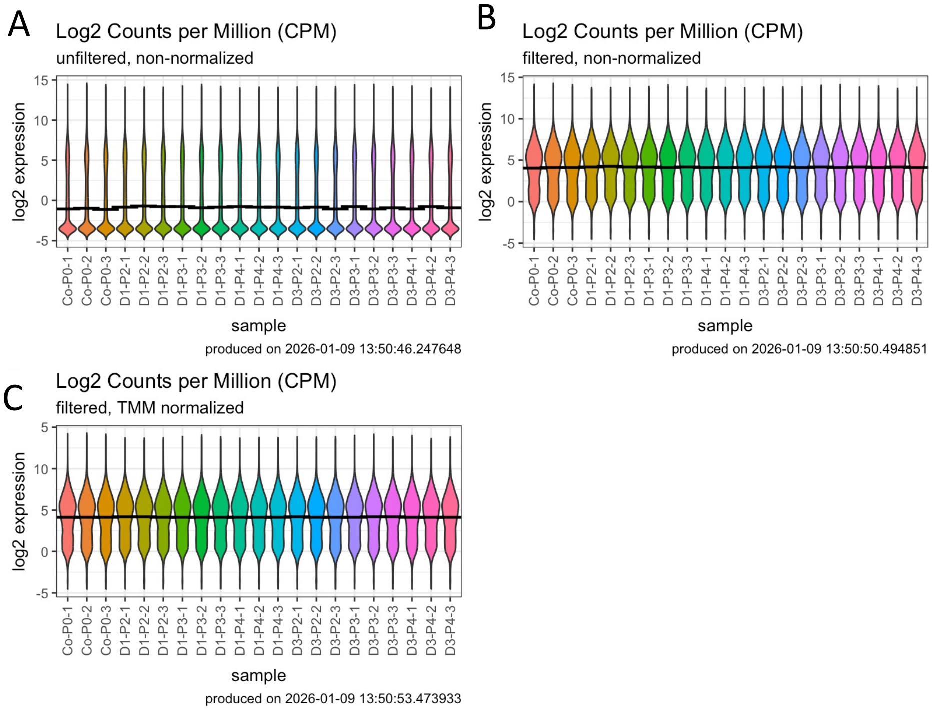


**Figure S2.** $Log_2$ transformation, normalization, and filtering of bulk RNA sequencing data for all samples from *in vitro* PBMC exposure to DENV-1 (45AZ5) and DENV-3 (CH53489). **A**) $Log_2$ CPM unfiltered, non-normalized. **B)** $Log_2$ CPM filtered, non-normalized. **C)** $log_2$ CPM filtered, normalized.

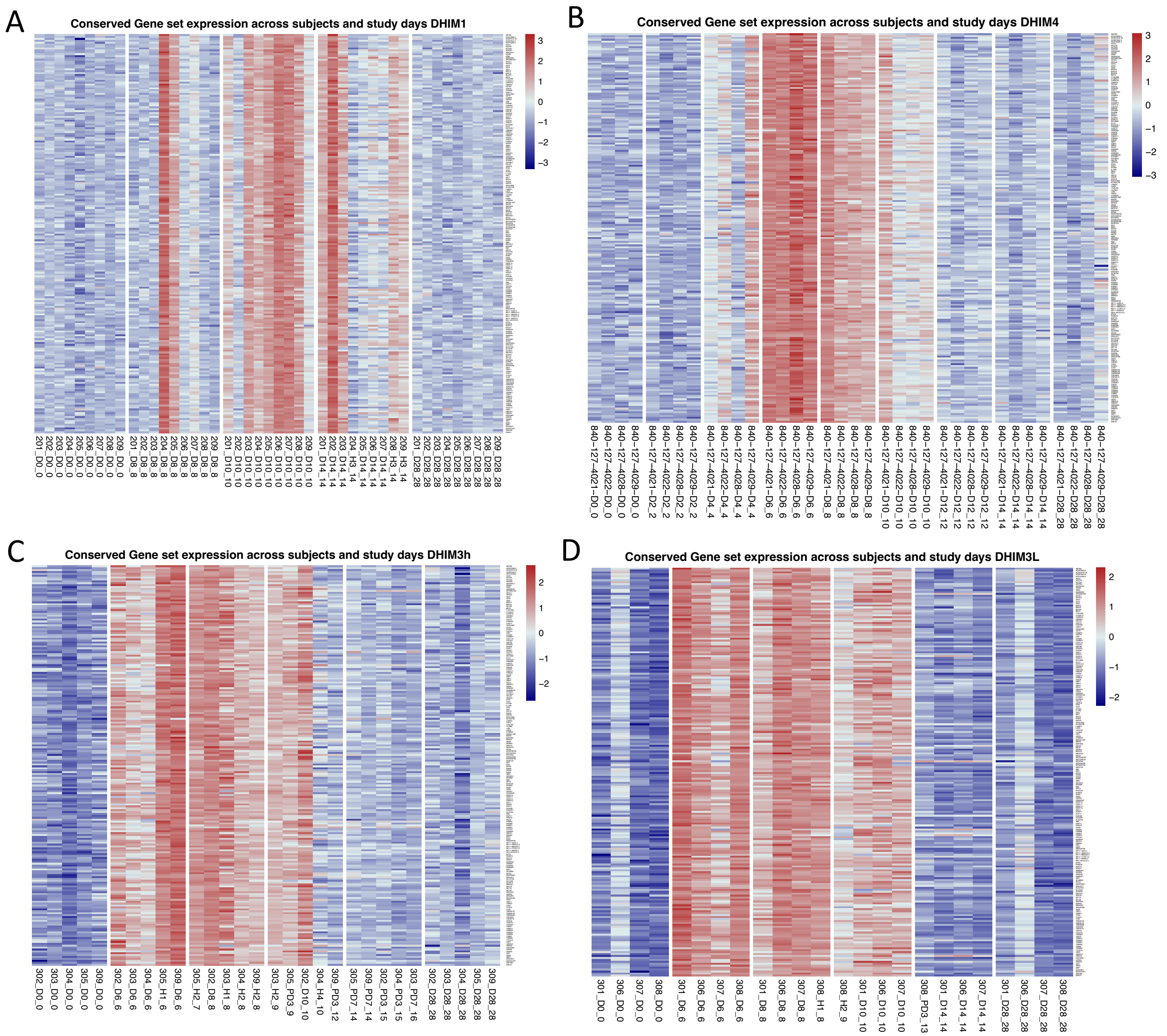


**Figure S3.** Heatmap of conserved upregulated gene set (n=210) expression across study days and cohorts. **A)** DHIM-1 cohort **B)** DHIM-4 cohort **C)** DHIM-3$^{high}$ cohort **D)** DHIM-3$^{low}$ cohort.

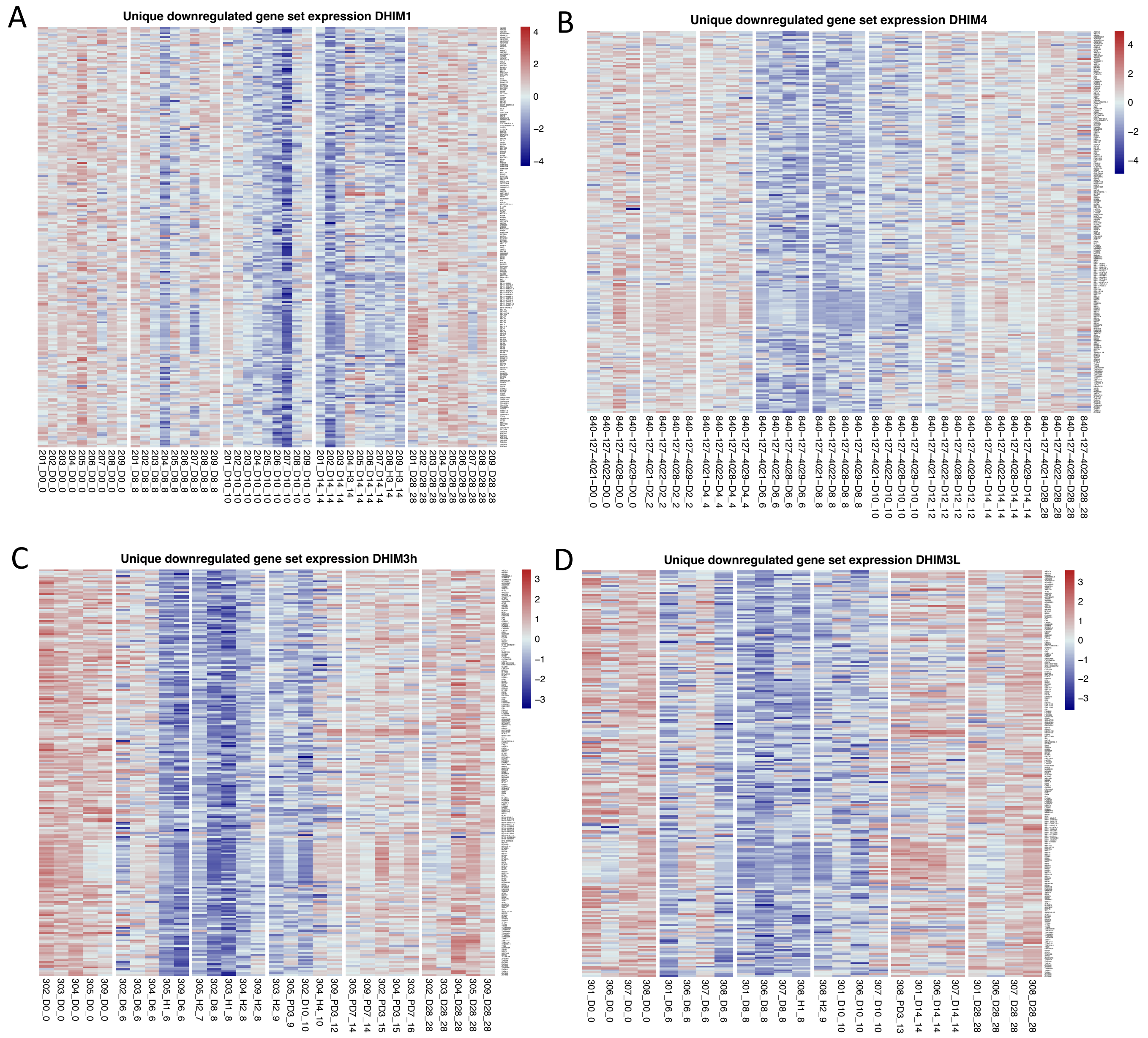


**Figure S4.** Heatmap of DHIM-3$^{high}$ unique downregulated gene set (n=224) expression across study days and cohorts. **A)** DHIM-1 cohort **B)** DHIM-4 cohort **C)** DHIM-3$^{high}$ cohort **D)** DHIM-3$^{low}$ cohort.

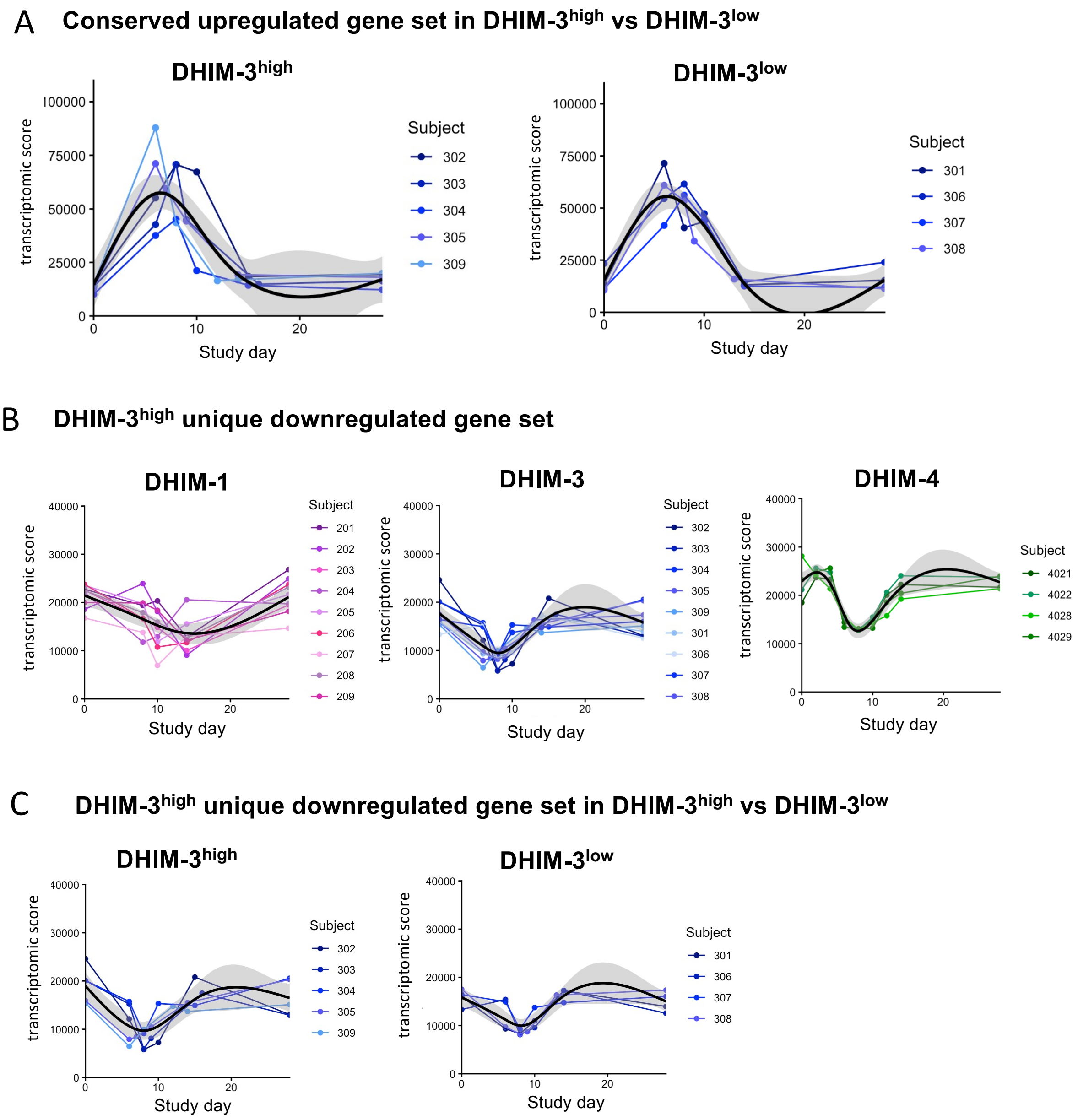


**Figure S5.** Transcriptomic score expression in DHIM-3$^{high}$ and DHIM-3$^{low}$ groups across study days. **A)** Conserved upregulated gene set (n=210) in DHIM-3$^{high}$ vs DHIM-3$^{low}$ **B)** DHIM-3$^{high}$ unique downregulated gene set (n=224). **C)** DHIM-3$^{high}$ unique downregulated gene set (n=224)in DHIM-3$^{high}$ vs DHIM-3$^{low}$ participants. Dots represent individual samples. A spline-based smoothing curve (black line) illustrates population-level trend between X and Y, with the grey region indicating the 95% confidence interval.

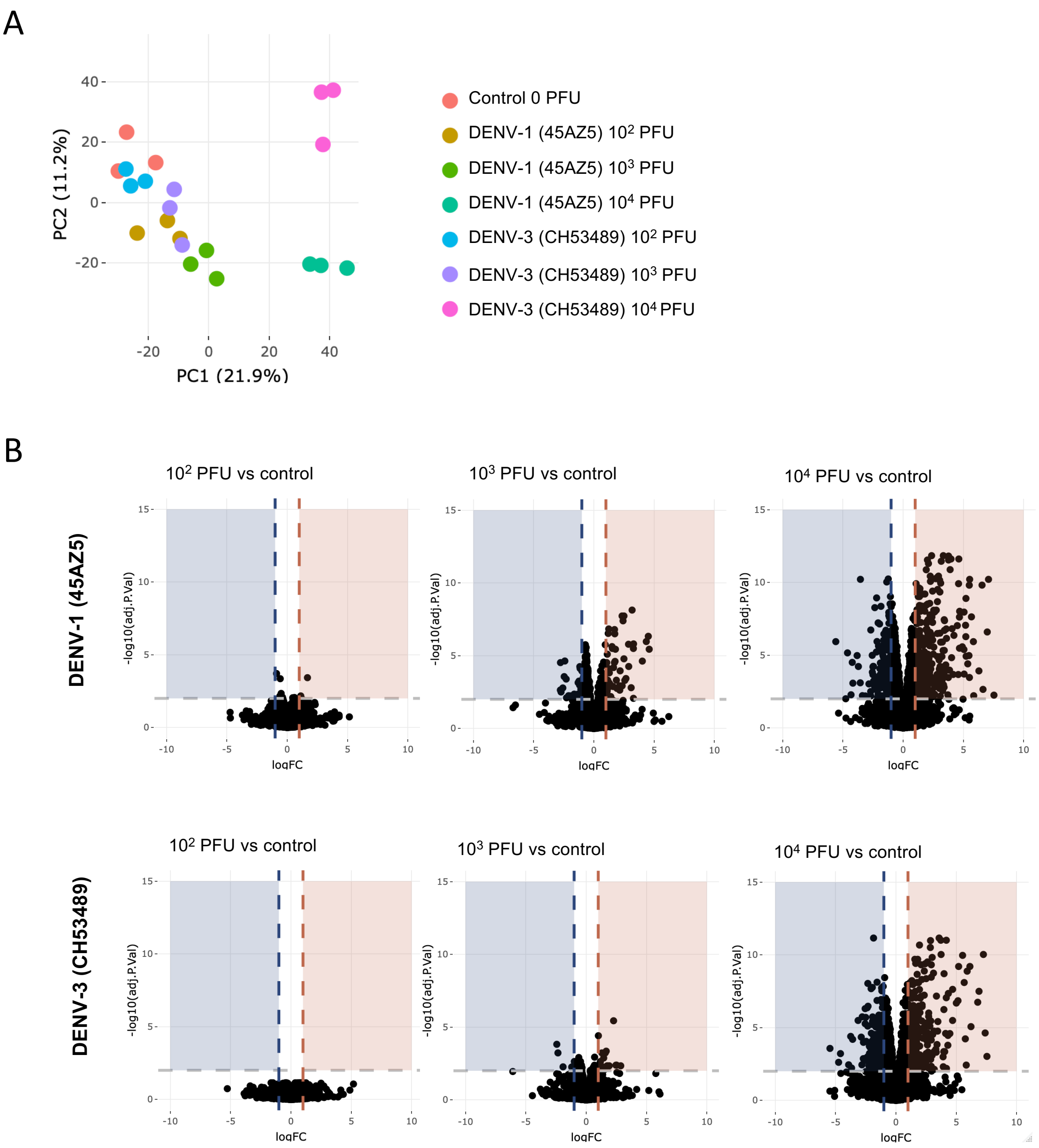


**Figure S6.** PCA analysis and volcano plots of DEG analysis for *in vitro* PBMC DENV-1 (45AZ5) and DENV-3 (CH53489) challenge strain titration. **A)** PCA analysis with individual samples labelled according to virus and PFU. **B)** Volcano plots indicating differentially expressed genes at $10^2$, $10^3$, or $10^4$ PFU compared to mock control.
